# Seasonal and hourly diversity patterns of anthropophagous female mosquito species in a semi-conserved area at the southern Mexico

**DOI:** 10.1101/2024.06.25.600586

**Authors:** Julio César Canales-Delgadillo, Nallely Vázquez-Pérez, Vicente Viveros-Santos, Rosela Pérez-Ceballos, José Gilberto Cardoso-Mohedano, Arturo Zaldívar-Jiménez, Omar Celis-Hernández, Alejandro Gómez-Ponce, Martín Merino-Ibarra

## Abstract

Mosquitoes are the most dangerous organisms on Earth because they spread causal agents of diseases. However, mosquito populations need to be better known in coastal areas of the Yucatan Peninsula to understand and prevent the spread of diseases effectively. To increase the knowledge about the mosquito community in the southern Gulf of Mexico region, we determined the diversity of mosquito species in Isla del Carmen Campeche. We trapped adult mosquitoes using buccal aspirators on monthly surveys (September 2019 to December 2020) in mangrove and low-semideciduous forest patches in three climate seasons. The sampling sessions consisted of 60 minutes of trapping performed every four hours (at 09, 13, 17, 21, 01, and 05 hrs) until a 24-hour cycle was completed. The trapped individuals were identified at the species level. We calculated Hill numbers using incidence data for each season and sampling hour. Abundances were compared through Kruskal-Wallis tests. A literature review determined the diseases associated with the species found. We collected 21,424 mosquito individuals from 11 genera, 26 species and four morphospecies. Seasonally, greater mosquito abundance and richness (*n* = 26) occurred during the norte season (*χ2* = 7.23, df = 2, *p* = 0.026) and between the 09:00 and 13:00 hrs (*χ2* = 15.25, df = 5, *p* = 0.009). Many species in this study are reported as disease vectors of medical and veterinary relevance. **Conclusions:** Isla del Carmen contributes to the Yucatan Peninsula’s mosquito diversity, and several of its species are vectors of pathogens to humans and wildlife.

## Introduction

The southeastern Gulf of Mexico (SGoM) is a region that stands out for its marine and terrestrial biodiversity and vast forested areas of ecological importance [1]. The wetlands, jungles, coastal areas, mangroves, and estuaries in this area provide wildlife shelter, food, and breeding sites [2]. Moreover, along the SGoM, numerous stopover sites are used for migratory birds as passage or rest areas during the fall and spring [3] because the Mississippi and Atlantic flyways converge over the Yucatan Peninsula.

Unlike mammals, birds are less susceptible to zoonotic agents [4,5]. However, due to their capacity to fly and migrate, they effectively distribute zoonoses of importance to humans and wildlife [6]. They act as natural hosts, reservoirs, or bridge hosts [7] to zoonotic agents that are transmitted by arthropod insects such as ticks and mosquitoes [8–10].

According with the last catalog of mosquitoes 3,570 species have been recorded worldwide (Wilkerson et al. 2021), and approximately 6% are present in Mexico [11]. Mosquitoes are widely regarded as among the most dangerous organisms on Earth, mainly because they can transmit pathogens that cause diseases, spreading them between humans or through zoonoses from animals to humans [12]. These tiny insects are responsible for transmitting a wide range of diseases such as malaria, dengue fever, chikungunya, yellow fever, and Zika virus [13]. Mosquito-borne diseases affect millions of people annually, particularly in tropical regions. Therefore, understanding the diversity of mosquito species in these regions is crucial for controlling and preventing the spread of vector-borne diseases (VBDs) [14].

VBDs are caused by pathogens transmitted to humans and domestic and wild animals through the bite of infected mosquitoes [13,14]. The transmission of VBDs is influenced by various factors, including the abundance and diversity of mosquito species [15], the density of human and animal populations [16], and environmental factors such as climate and land use change [17–20].

Southern Mexico is a tropical hotspot area for mosquito-borne diseases [21]. The region is characterized by high temperatures, high humidity, and abundant water bodies, creating an ideal mosquito breeding environment [22]. As a result, several mosquito species incriminated in transmitting VBDs inhabit this region [21,23]. For example, Campeche, located in the Yucatan Peninsula (YP), is home to numerous mosquito species that are incriminated in transmitting various VBDs [24]. According with Talaga et al. (2023), 90 species are recognized as occurring in YP. However, additional studies on the species diversity of mosquitoes are needed because there are still significant gaps in the knowledge on the ecology and distribution of these insects. For instance, *Aedes* (*Stegomyia*) *aegypti* Linnaeus, 1762, and *Anopheles* (*Nyssorhynchus*) *albimanus* Wiedemann, 1820, are well-known on the YP. Nevertheless, other mosquito species may also be relevant disease vectors [25], and their distribution and abundance have not yet been studied. Therefore, studies contributing to understanding the region’s full range of mosquito species are crucial for help designing effective disease control and prevention strategies.

Isla del Carmen is a small island off the coast of Campeche, Mexico. The island environment is characterized by humid tropical weather, warm temperatures and high levels of rainfall throughout the year. The island is known for its mangrove ecosystem [26,27], which provides an ideal breeding ground for mosquitoes because the mangrove trees and their associated vegetation offer many microhabitats for mosquitoes to breed, feed, and rest [28]. Despite the importance of the Isla del Carmen environment in supporting mosquito populations, knowledge about its mosquito species diversity could be clearer. Only a few studies have focused on Campeche mosquito species. For example, Bond et al. [29] focused on species of medical importance distributed in areas of low semievergreen forests and mangroves in continental areas of Campeche, which is far from Isla del Carmen. Others have studied mosquitoes inhabiting primary and secondary low semievergreen forest habitats and fruit tree plantations adjacent to riverine ecosystems [24]. However, mosquito species diversity and abundance on the island are not well-known. Moreover, there is only one published work from a survey conducted between 1996 and 2006, where the authors reported 14 mosquito species of medical relevance for Isla del Carmen [30]. Therefore, we aimed to determine the diversity of anthropophagous female mosquito species in Isla del Carmen seasonally and within 24-hour cycles to increase the knowledge about this group of insects in the coastal areas of the southern Mexico region. In addition, because of the environmental characteristics of our study area, we expected the species richness and diversity to be similar to those reported for other areas of the YP. Lastly, our work will provide baseline data for future assessments of how mosquito populations and their associated pathogens could change over time as climate and ecosystem modifications occur.

## Materials and Methods

### Study area

Isla del Carmen is a sandbar island located southwest of the Mexican state of Campeche, within the YP (Fig. 1). The climate is warm subhumid with minimal and maximal average temperatures ranging from 22°C to 32°C and approximately 1155 mm of rainfall annually. Around the year, the relative moisture content is approximately 74%. The dominant vegetation types are mangroves and semideciduous forests.

**Figure 1.**
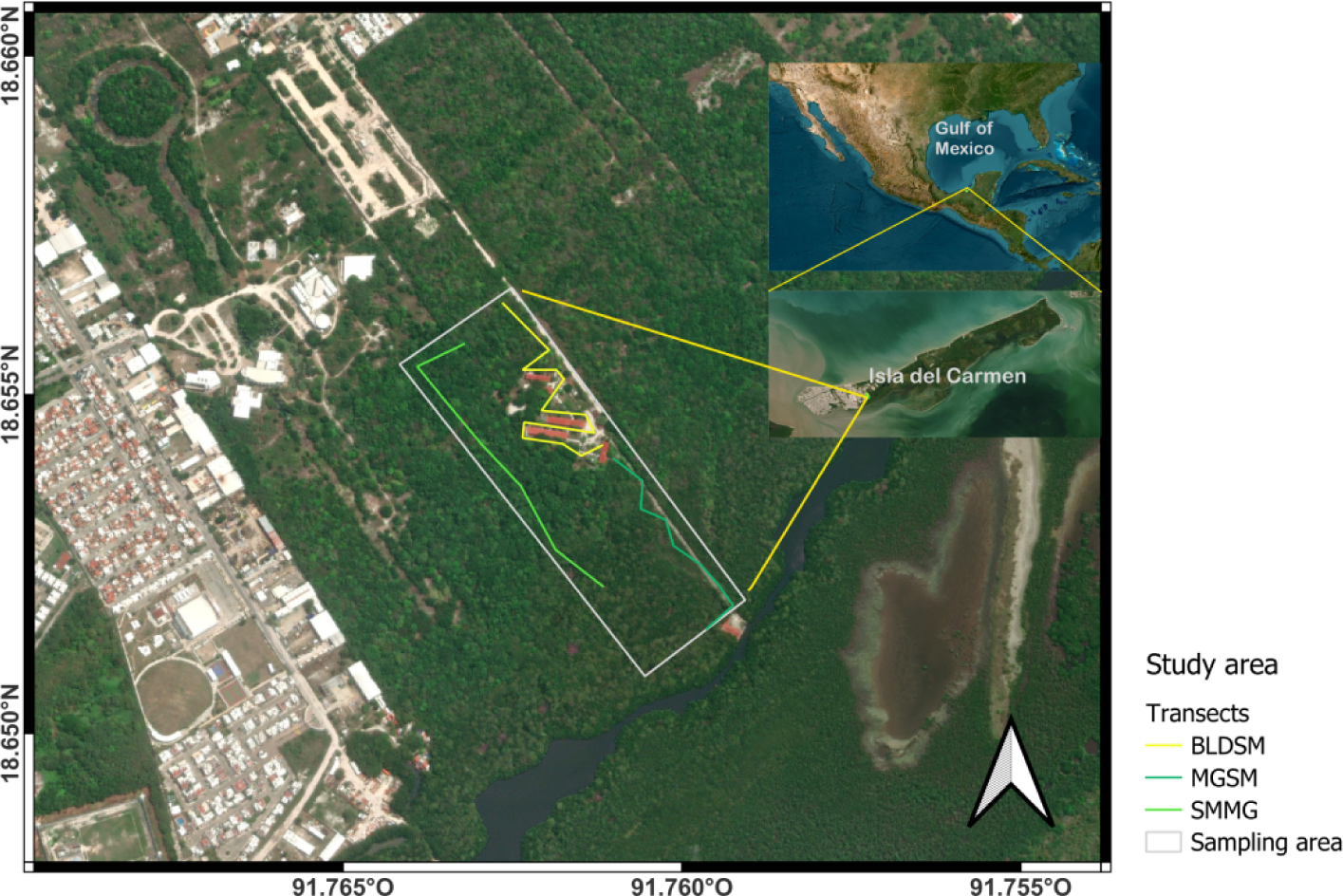
Study are and sampling transects. The transects were separated by dominant vegetation: buildings and semideciduos forest (BLDSM), mangorove and semideciduos forest (MGSM), semideciduos forest and mangrove (SMMG).

The sampling site (18° 39’ 13’’ N, -91° 45’ 41’’W, Fig. 1) is a ten-hectare area, modified by the presence of buildings for academic research and embedded in a changing landscape of growing human settlements located at an approximate average distance of 546 m. Although this human influence is evident, within the sampling area four species of mangrove trees (*Rizophora mangle* L.,*, Laguncularia racemose* (L.) C. F. Gaertn*, Avicennia germinans* (L.) L.*, Conocarpus erectus* L.), form an ecotone with a vegetation community composed of *Metopium brownie* (Jacq.) Urb.*, Bursera simaruba* (L.) Sarg. 1890*, Sabal Mexicana* Mart.*, Lonchocarpus hondurensis* Benth., and *Cedrela odorata* L., among other plant species typical of tropical semideciduous forests. Therefore, the association between mangroves and the tropical semideciduous forests within our study site represents the typical vegetation community of disturbed and natural habitats in Isla del Carmen. Regional climate seasons are three: nortes or cold front season (NS) from October to February; dry season (DS) from March to May; and rainy season (RS) from June to September [31,32].

### Sampling

To assess the anthropophagous mosquito species diversity, we carried out monthly samplings, from September 2019 to December 2020 (permit number: SGPA/DGVS/0408019). The sampling included three collectors that used pooters (oral aspirators) to trap live-flying adult mosquitoes on three sampling transects with vegetation dominated by mangrove, semideciduos forest, or semideciduos forest modified by buidings (Fig. 1), through the human landing collection (HLC) method [ECDC and EFSA 33]. The HLC technique is one of the oldest methods to catch mosquitoes and has been widely used for its simplicity in collecting anthropophagous host-seeking females [ECDC and EFSA 33,34]. Before starting the sampling, collectors were informed about the risks related to vector-borne diseases and mosquito biting within the region of the study site. Once collectors agreed with working conditions, the sampling began. To minimize the infection risk from mosquito bites, collectors used denim clothes and latex gloves to protect as much skin as possible. During the study period, collectors did not show any infection symptoms.

During the sampling, we covered 16 24-hour cycles consisting of six 60-mins trapping sessions per cycle separated by three hours each: in the mornings at 05 and 09 hrs., in the afternoons at 13 and 17 hrs., and in the night at 21 and 01 hrs., (Montarsi et al. 2015; Santos et al. 2020). In addition, for each sampling session one collector walked on one of the three sampling transects (Fig. 1), to trap mosquito females seeking for a host. Each collector was equipped only with an LED head lamp for secure walking after sunset, one 1-L plastic container for mosquito storage, and a pooter composed of a 30 cm glass tube connected to an 85 cm latex hosepipe with filters at each extreme of the pipe. No UV light or other attractors were used for sampling. Thus, the sampling effort was six hours per transect which summed 18 hours per sampling, for a total of 288 hours of sampling time along the three climate seasons in our study area. Mosquitoes were trapped mainly from the collectors’ bodies, but when possible, also from bodies of domestic dogs. The trapped individuals were stored in 1-L plastic containers until they were transported to the laboratory for sacrifice via thermal shock (-20°C) for 5 to 10 mins. All the trapped individuals were inspected under a stereoscope (Stemi 305, Carl Zeiss Oberkochen, Germany) for taxonomical identification using specialized keys such as Carpenter and LaCasse (1955), Díaz-Nájera [35], Arnell (1976), Clark-Gil and Darsie [36], and Wilkerson et al. [37]. Once separated by species, all the mosquitoes were counted, some specimens were prepared and mounted for a reference collection which was deposited at the Biodiversity and Conservation Genetics laboratory at the Institute of Sea Sciences and Limnology El Carmen UNAM.

### Data analyses

For diversity analyses, the data were grouped according to the seasonal pattern occurring in our study area (DS, RS, NS) and the 24-hours sampling cycle.

To evaluate the sampling coverage during different seasons and at different times of the 24-hour cycle, we generated accumulation diversity curves. After the conversion of the abundance data to incidence data, we generated three estimates of Hill numbers of order *q*: 1) species richness (*q* = 0); 2) Shannon diversity, as the exponential of Shannon entropy or common species (*q* = 1); and 3) Simpson diversity, calculated as the inverse of Simpson concentration or dominant species (*q* = 2), as implemented in the R package iNEXT [38,39].

The seasonal and 24-hour abundances were compared using nonparametric Kruskal-Wallis’ test. To better comprehend the contribution of the Isla del Carmen mosquito species to regional diversity, we compared our data with diversity reports in other areas of the YP. We compiled published works with available abundance data [24,29,40–45], to estimate species diversity and the relative abundance of each reported species in each state/location as described above. All analyses were performed in R version 4.1.1 [46].

## Results

### Species richness and abundance

During the sampling period, we collected 21,424 mosquito individuals, from which 98.60% (*n* = 21,124) were adult females, and 1.40% (*n* = 300) were adult males, divided into two subfamilies (Culicinae and Anophelinae), 11 genera, 26 species and four morphospecies (Table 1). About 99% of the captured females showed empty abdomen, 147 individuals were completely engorged, and 64 were partially engorged, showing dark blood within the abdomen, meaning that blood meals were taken less than six hours or between six and 18 hours before being trapped, respectively. No gravid females were collected during the samplings. Collected males belonged to the species *Ae.* (*Ochlerotatus*) *taeniorhynchus* Wiedemann, 1821 (*n* = 263), *Culex* (*Culex*) *interrogator* Dyar & Knab, 1906 (*n* = 15), *Psorophora* (*Janthinosoma*) *ferox* von Humbolt, 1819 (*n* = 10), *An.* (*Anopheles*) *crucians* Wiedemann, 1828 (*n* = 7), and *Ae.* (*Ochlerotatus*) *angustivittatus* Dyar & Knab, 1907 (*n* = 5). Most of the females were caught on the collectors’ bodies. However some catches (< 1%) were obtained from domestic dogs’ bodies (*Ae. taeniorhynchus*, *An. crucians*). Only males were caught mainly from buildings’ windows and walls (*Ae. taeniorhynchus*, *An. crucians*, *Cx. interrogator*), and from plant branches and leafs (*Ps. ferox*, *Ae. angustivittatus*).

**Table 1.**
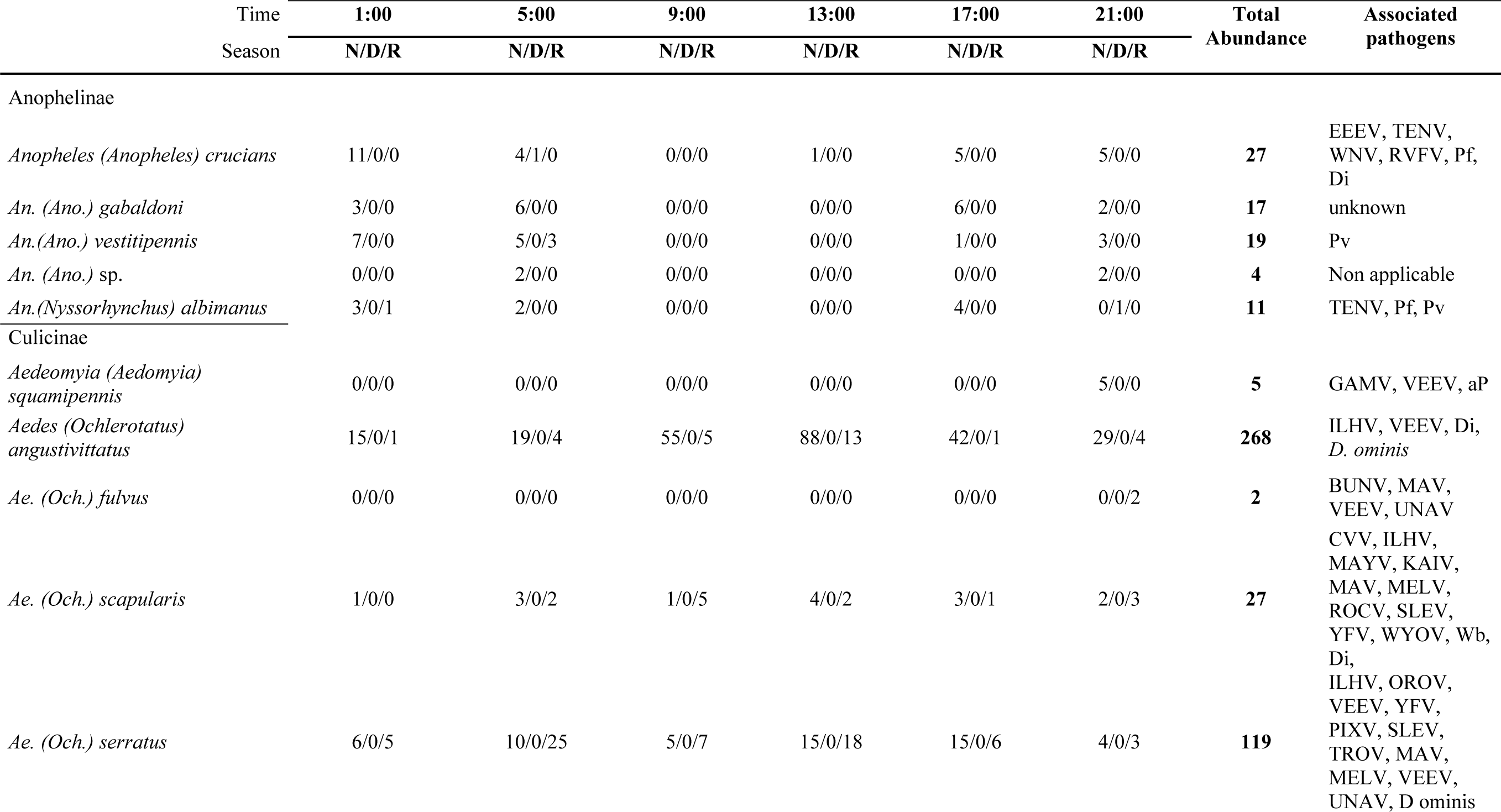
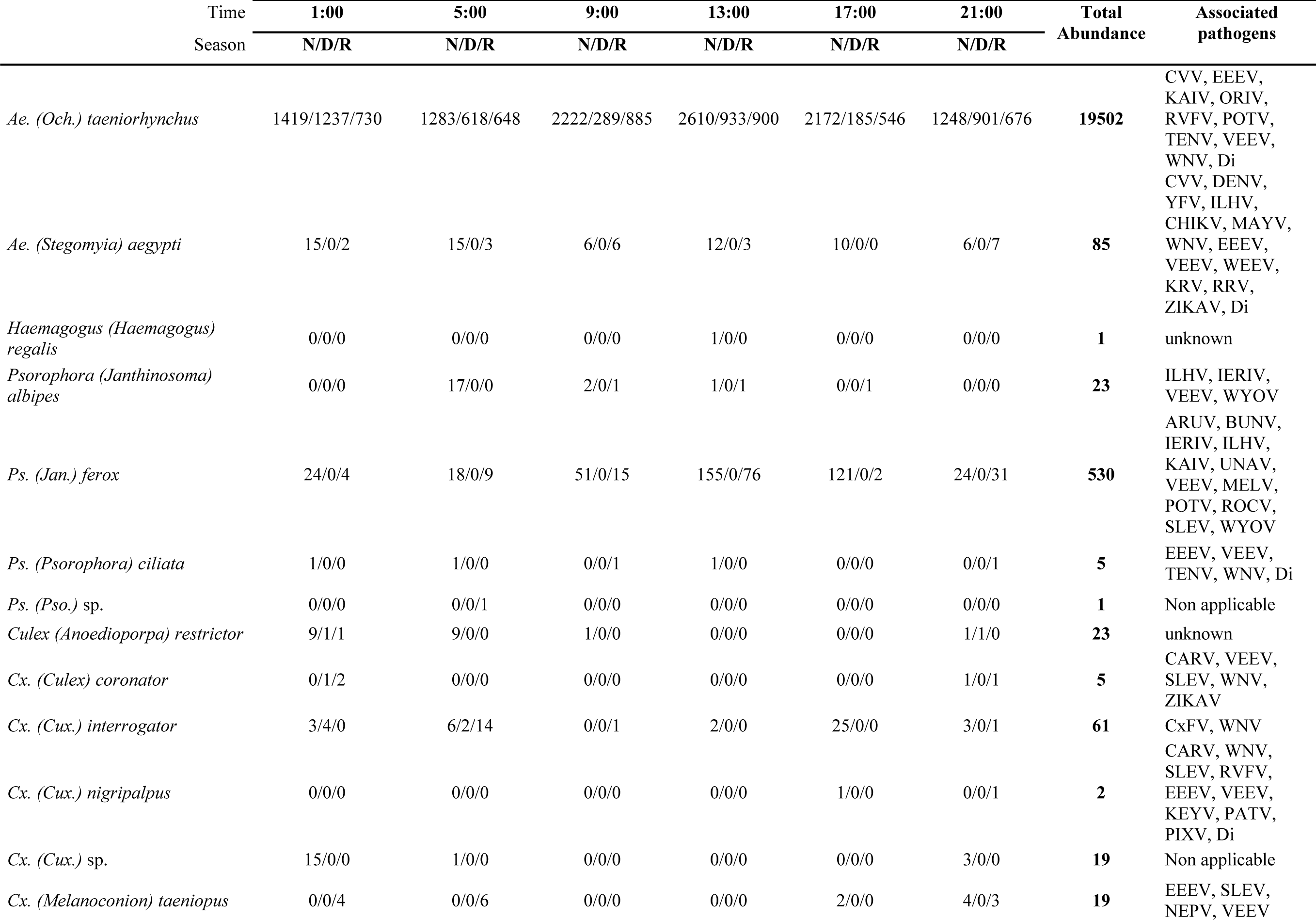
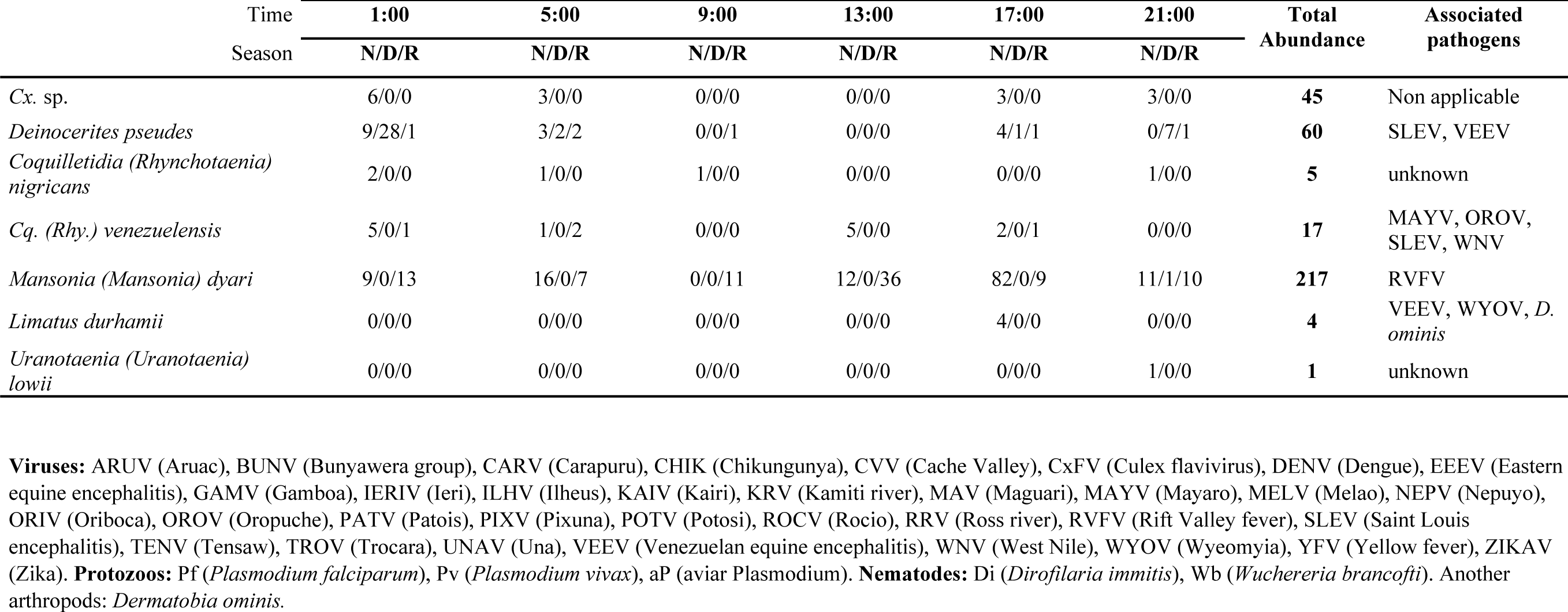
Species richness and abundance of anthropophagous female mosquitoes in Isla del Carmen during a 24-hour cycle and three sampling seasons: Norte (N), dry (D), and rainy (R). The medical and veterinary relevance of mosquito species was determined by the associated pathogens reported in the literature.

All the genera recorded in our study were present in NS, while in DS, less than the 50% of the recorded genera were present (*Anopheles* Meigen*, Aedes* Meigen*, Culex* Linnaeus*, Deinocerites* Theobald, and *Mansonia* Blanchard). However, *An. albimanus, Ae. taeniorhynchus, Cx.* (*Anoedioporpa*) *restrictor* Dyar & Knab, 1906, *Cx.* (*Culex*) *coronator* Dyar & Knab, 1906, *Cx. interrogator*, *De. pseudes* Dyar & Knab, 1909, and *Ma.* (*Mansonia*) *dyari* Belkin, Heinemann & Page, 1970, were recorded in all sampling seasons.

The highest species richness was recorded during the NS (*n* = 26), while it was lower in the RS and DS (*n* = 20, and *n* = 8, respectively, Table 2). The black salt marsh mosquito (*Ae. taeniorhynchus*) was the most abundant and dominant species at any time during the 24- hour cycle, sampling seasons and vegetation type, representing 92.3% of our sample (Fig. 2).

**Figure 2.**
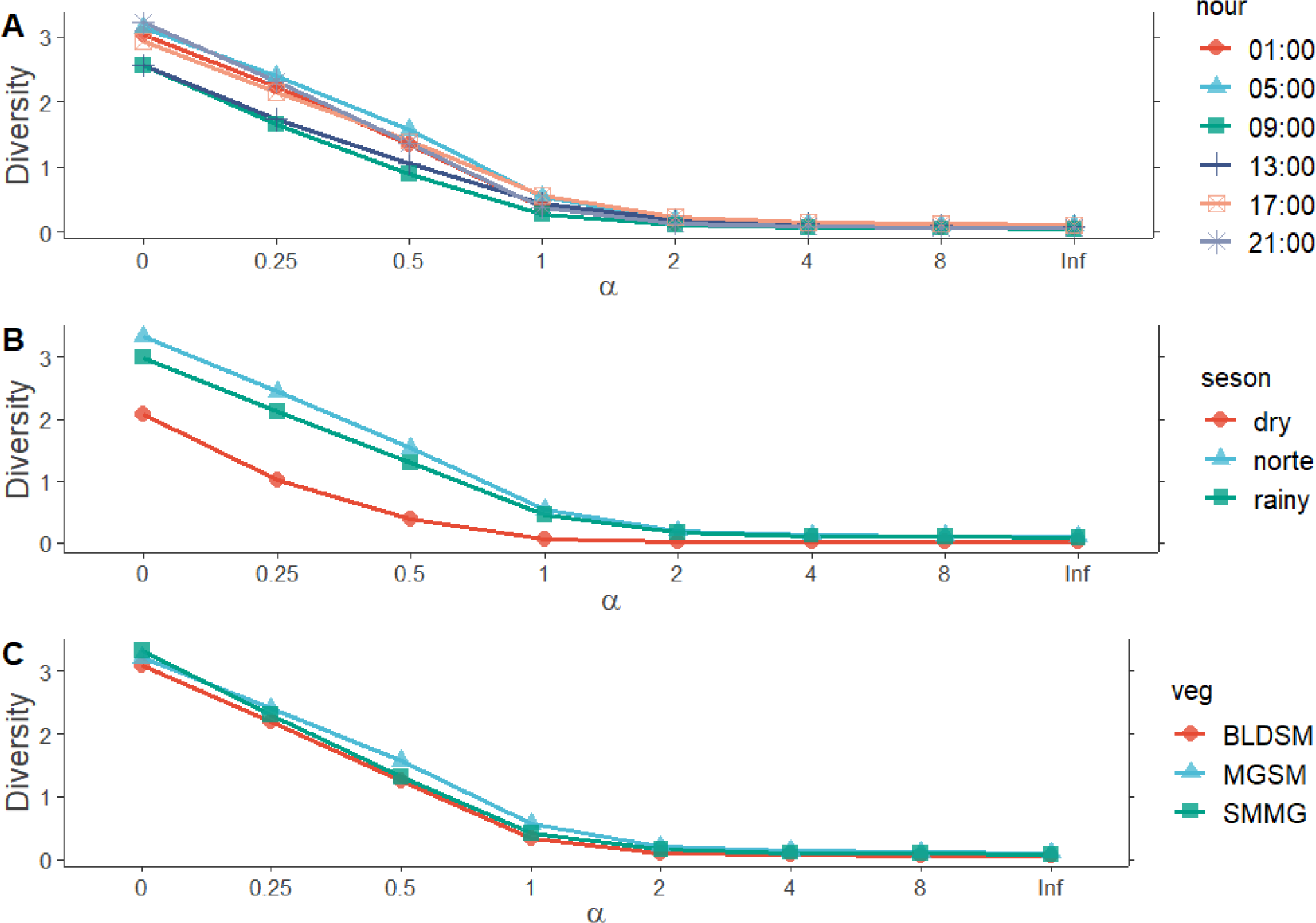
Renyi diversity profiles of female mosquitoes in Isla del Carmen by hourly activity (A), season (B), and vegetation type (C). In the plots, α = 0 is the species richness, α = 1 the Shannon-Weiner diversity index, α = 2 the Simpson diversity index, and *Inf* is the Berger-Parker dominance index.

**Table 2.**
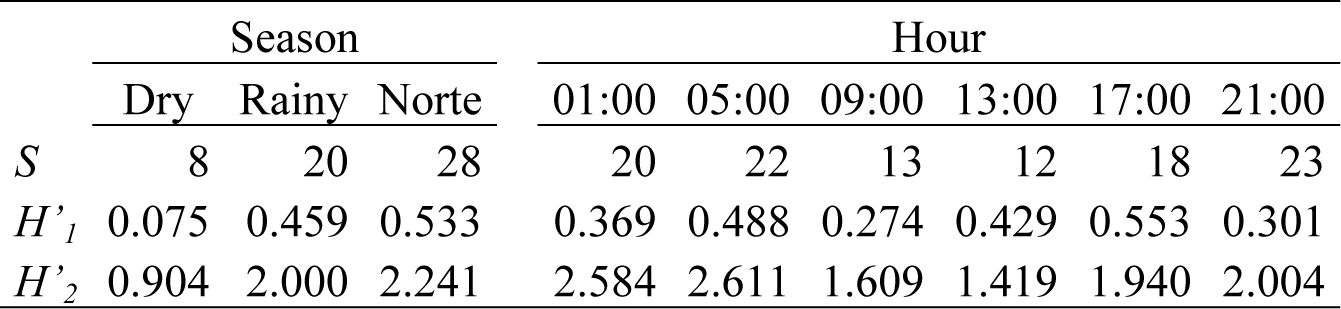
Seasonal and hourly mosquito species richness (*S*) and diversity (*H’*) in Isla del Carmen. To show the effect of an eudominant species on the local community, we compared the data including (*H’_1_*) and excluding (*H’_2_*) *Ae. taeniorhynchus*.

|  | Season |  |  | Hour |  |  |  |  |  |
| --- | --- | --- | --- | --- | --- | --- | --- | --- | --- |
|  | Dry | Rainy | Norte | 01:00 | 05:00 | 09:00 | 13:00 | 17:00 | 21:00 |
| $S$ | 8 | 20 | 28 | 20 | 22 | 13 | 12 | 18 | 23 |
| $H'_1$ | 0.075 | 0.459 | 0.533 | 0.369 | 0.488 | 0.274 | 0.429 | 0.553 | 0.301 |
| $H'_2$ | 0.904 | 2.000 | 2.241 | 2.584 | 2.611 | 1.609 | 1.419 | 1.940 | 2.004 |

The genus *Aedes* was the best represented (94.6%), while genera such as *Aedeomyia* Theobald*, Deinocerites, Limatus* Theobald, and *Uranotaenia* Lynch Arribálzaga, typically had low relative abundances (0.023, 0.284, 0.018, and 0.004%, respectively). Seasonally, a greater number of mosquitoes occurred during the NS. Even when excluding the most abundant species (*Ae. taeniorhynchus*) from the data, the NS had a greater mosquito abundance (Kruskal−Wallis test *χ^2^* = 7.23, df = 2, *p* = 0.026). Mosquito abundance also varied with mosquito timing between the 09:00 and 13:00 hrs (Kruskal−Wallis test *χ^2^* = 15.25, df = 5, *p* = 0.009).

### Diversity patterns

The diversity analysis revealed values greater than 90% sample coverage in the NS and RS (94.71 and 92.1%, respectively). A lower sample coverage was estimated for the DS (81.48%). In addition, we had sample coverage greater than 80% during each sampling hour (except at 17:00, where sample coverage was 75.5%), indicating that our sampling effort was sufficient to determine the female mosquito species richness in our study area.

The Hill numbers showed that the season with the highest diversity of culicids was the NS. We observed that common (*q* = 1) and dominant species (*q* = 2) were significantly different for all the seasons (because no confidence intervals overlapped). However, for species richness (*q* = 0), the differences between RS and NS were not clear, even when we extrapolated our sampling effort twice (Fig. 3), meaning that these two seasons had similar diversity patterns. When we considered the diversity of each season separately, we found that diversity was on average 3.8 and 1.4 times greater in the NS than in the DS and RS seasons, respectively. Moreover, the RS was on average 2.6 times more diverse than DS. Finally, the estimated number of undetected species for each season was NS = 7.5, DS = 3.3, and RS = 7.5 (Fig. 3).

**Figure 3.**
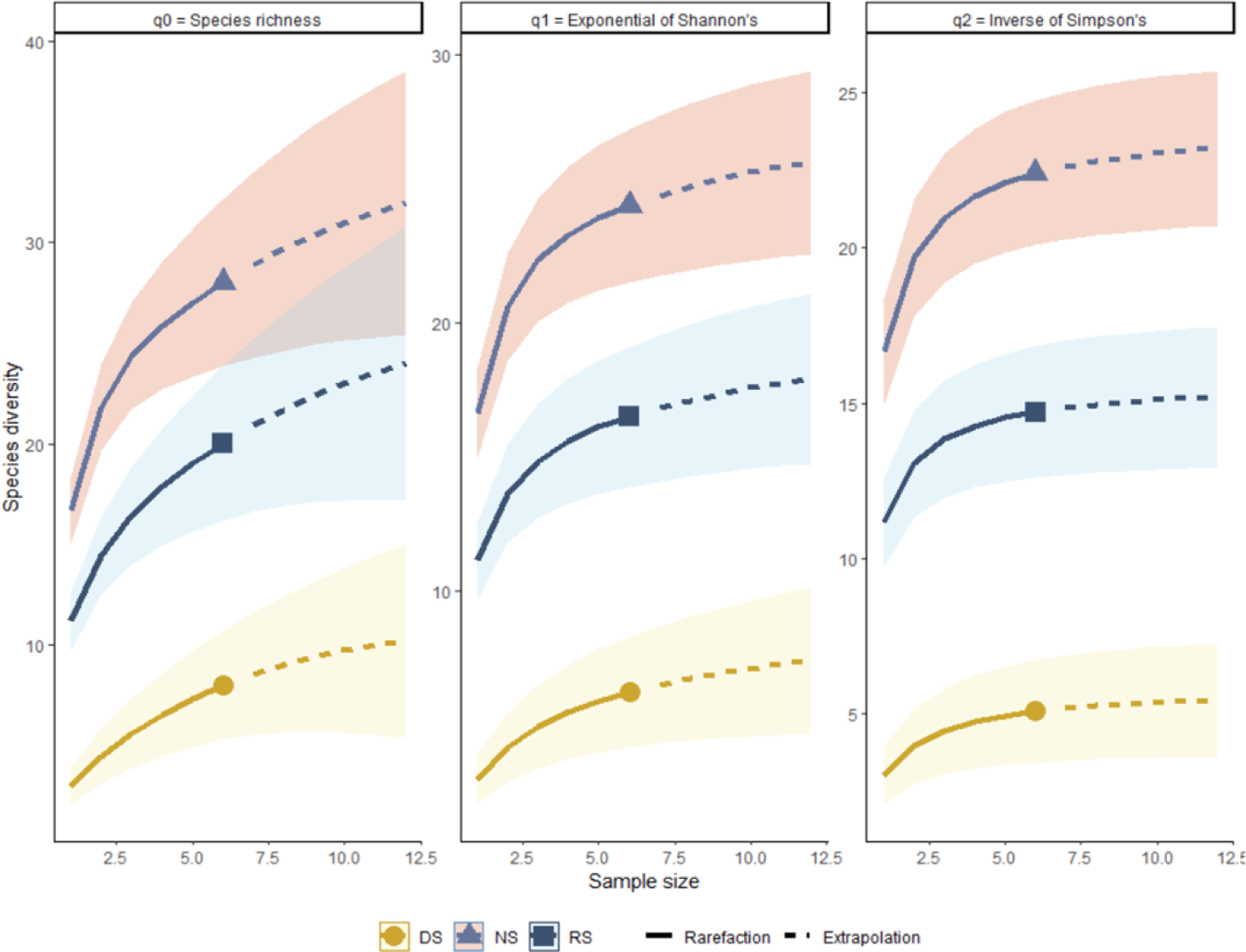
Estimated Hill numbers for species richness (*q* = 0), equally common species (*q* = 1), and the effective number of dominant species (*q* = 2) during the three sampling seasons. No overlap of the confidence intervals (shaded areas) indicates significant differences.

The rarefaction analyses showed that the Hill numbers differed significantly among seasons (Figure 4A), indicating that the mosquito community of Isla del Carmen has a varying dynamic throughout the year, except for the dominant *Ae. taeniorhynchus*, which is highly abundant in all seasons. The rest of the species are likely affected by yearly environmental changes. In addition, during the study period, the highest species richness occurred from 17:00 to 05:00 hours, while the lowest number of species occurred between 09:00 and 13:00 (Figure 4B).

**Figure 4.**
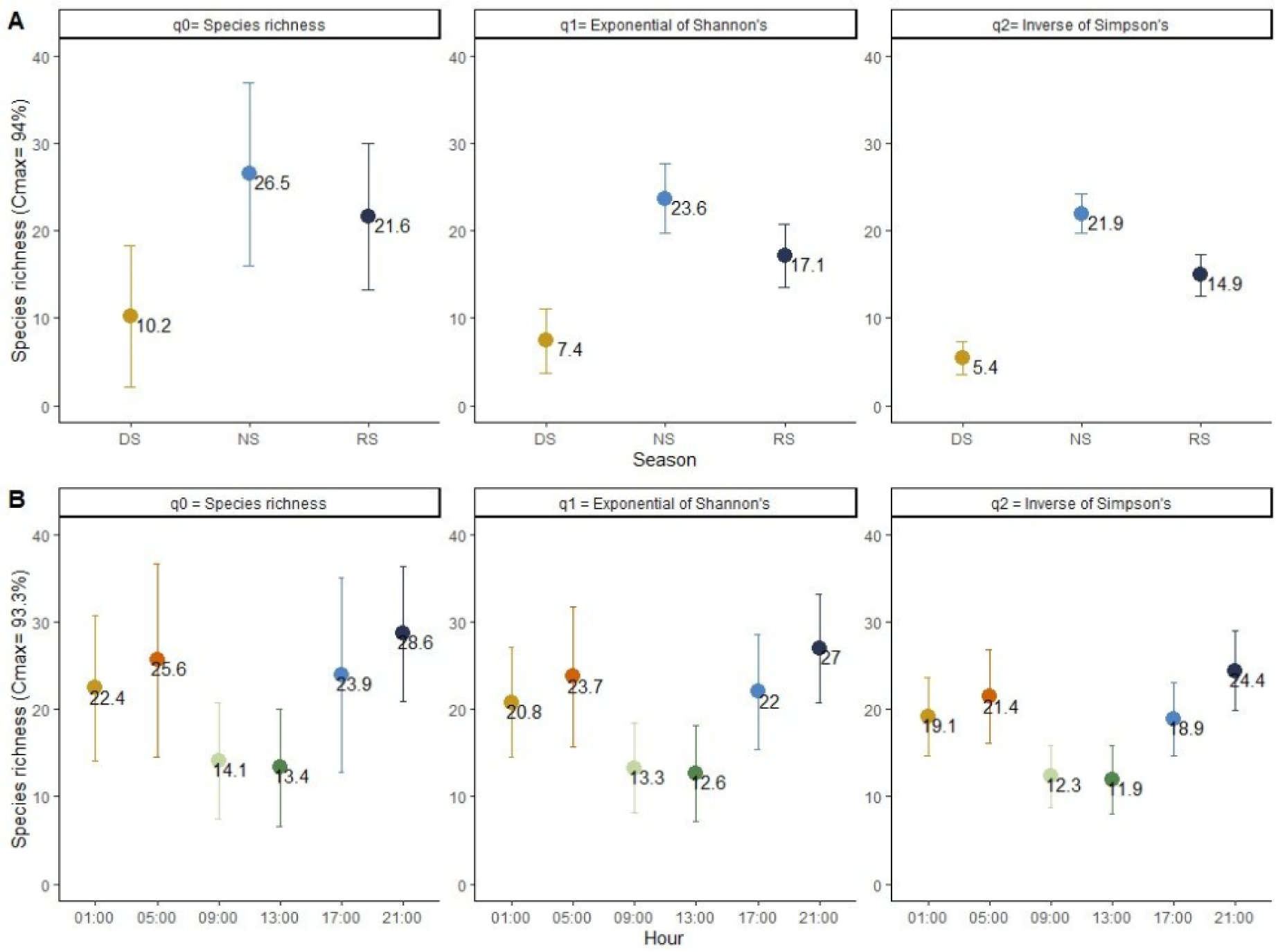
Rarefied estimates of species richness and diversity patterns in three sampling seasons (A) and six different sampling hours (B) are shown. Estimations are based on the sample maximal coverage (Cmax). Vertical bars represent the 95% lower and upper confidence limits.

According to the reviewed literature, Isla del Carmen is among the places with the greatest species richness on the YP, together with Celestún and Calakmul. However, it showed a diversity value similar to that of the urban areas of Merida city. But, when the eudominant *Ae. taeniorhynchus* was excluded from the dataset, the diversity of the mosquito community in Isla del Carmen was greater than the estimated value for all other locations (Table 3), like the pattern observed locally (see Table 2).

**Table 3:** Female mosquito species richness (*S_r_*), and diversity of Isla del Carmen as compared with other localities in the Yucatan Peninsula. Parameters were estimated from data of this study and earlier published works. The estimated Shannon-Wiener diversity index (*H’*) by locality included (*H’_1_*) and excluded (*H’_2_*) *Ae. taeniorhynchus* data to show the effect of this eudominant species on the communities.

| Parameter | Isla del Carmen | Candelaria | Celestún | Mérida | Palizada | Campeche | Calakmul | Sian kaan |
| --- | --- | --- | --- | --- | --- | --- | --- | --- |
| $S_r$ | 26 | 23 | 33 | 19 | 11 | 6 | 30 | 26 |
| $H'_1$ | 0.447 | 1.607 | 1.629 | 0.751 | 1.377 | 0.752 | 2.089 | 1.761 |
| $H'_2$ | 2.296 | 1.659 | 2.144 | 1.320 | 1.095 | 0.752 | 2.074 | 1.572 |

## Discussion

For the first time, we report the 24-hour cycle dynamics of the Culicidae family during the different climate seasons occurring in Campeche, particularly within the Terminos Lagoon region [31]. By identifying and cataloging the mosquito species present in Isla del Carmen, our study provides crucial data on the local and regional mosquito fauna in the SGoM, particularly in the Yucatan Peninsula. This knowledge is foundational for understanding the pathobiology of mosquito-borne diseases, as different mosquito species can carry different pathogens [18,47].

The mosquito population and disease dynamics are affected by environmental factors such as precipitation, temperature, and relative humidity [48], by changes in mosquito mortality, and food and breeding site availability [49]. In our study, the NS was the season with the highest abundance (57.38%) and diversity values as compared to the DS and RS. This result is similar to that reported by Romero-Vega et al. [50], who reported that mosquito diversity is greater in the wet season than in the dry season.

According to Guerra-Santos and Khal [51], the NS and the RS in our study area have very similar monthly rainfall values. In addition to rainfall, the NS typically experiences temperature decreases because of the occurrence of cold fronts, which may favor bionomic conditions [52,53]. During the NS, the less abundant rainfall helps the immature mosquito phases have more favorable breeding site conditions for developing and increasing the number of emerging adults [54], in contrast with overflooding and water flow events that naturally eliminate the immature phases during the RS [55]. Roiz et al. [56] reported that rainfall accumulation is positively related to the abundance of several species of *Anopheles, Psorophora, Uranotaenia*, and subgenera *Ochlerotatus* Lynch Arribálzaga and *Aedes*, which need large breeding sites at ground level with vegetation and standing water [57–60]. Such conditions are more likely to occur during NS in our study area, increasing mosquito abundance. Likewise, species like *An. albimanus* and *Ae. taeniorhynchus, Cx. interrogator, De. pseudes* and *Ma. dyari* may breed in a broader range of environmental conditions since they were present throughout the study period. Furthermore, during the NS, cooler temperatures allow more species to experience activity during the day because of lower dehydration and rainfall probabilities. For example, Drakou et al. [49], reported that the optimal temperature for mosquito activity ranges from 15°C to 24°C, while above 28°C, the activity decreases. During the NS season, the average temperature in our study area was two Celsius degrees lower than in RS and DS seasons [61], which might favored increased mosquito abundance.

In our study, we did not find differences in the incidence of mosquitoes for most of the sampling hours, except for 09:00 and 13:00, when the number of species and the number of caught mosquitoes were lower. However, some of the recorded species, such as *An. crucians*, *An.* (*Ano.*) *gabaldoni* Vargas, 1941, and *An.* (*Ano.*) *vestitipennis* Dyar & Knab, 1906, were diurnally active, although the genus *Anopheles* is considered to have mainly nocturnal or crepuscular activity [62]. These species were inactive only between sunrise and 09:00 hours, a pattern reported for species of medical importance [63]. Conversely, *Ae. angustivittatus*, *Ae.* (*Och.*) *scapularis* (Rondani, 1848), *Ae.* (*Och.*) *serratus* (Theobald, 1901)*, Ae. aegypti*, and *Ps. ferox*, considered as diurnal species [64,65], were observed at dusk (21:00, 01:00 and 05:00). *Mansonia dyari* is also considered a nocturnal species [66,67], however during the RS and NS, this species was recorded at all sampling hours.

Throughout the study, the hour at which the highest species richness occurred was 21:00. Although at this time, most of the diurnal species (*Aedes, Psorophora, Haemagogus* Williston) exhibited a reduced incidence, some individuals could remain actively searching for a meal, co-occurring with the nocturnal species (*Anopheles, Coquillettidia* Dyar, and *Mansonia*) that exhibited increased activity [68]. However, the observed activity patterns could have been influenced by the presence of collectors [53], and additional studies are needed.

Although the sampling coverage was in the range of 81-94%, the analyses revealed a few undetected species (NS and RS = 7 species, DS = 3 species) that can be recorded by using additional catching methods and increasing the sampling effort to overcome the high presence of singletons that usually appear in entomological surveys [69].Isla del Carmen houses 44% of the mosquito species reported for the entire estate of Campeche [24,40,41], 35% of the species reported in Yucatan [44,45], and approximately 30% of the recorded species in Quintana Roo [40,41,43]. According to the recently published review of Talaga et al. [70], the species richness found in our study represents about 29% of the mosquito species recorded within the YP. Thus, the high level of species richness underscores the ecological importance of Isla del Carmen as a regional mosquito diversity hotspot, and its potential role in regional disease dynamics.

Although the species richness in Isla del Carmen was similar to that in other studied locations, its species diversity was lower than that in other coastal and inland areas of the YP because of the high numbers of *Ae. taeniorhynchus*, a eudominant species in our study area comprising approximately 92% of the captured adult females. As *Ae. taeniorhynchus* oviposits in brackish water [71], it is expected that the estuarine environment and the mangrove areas surrounding Isla del Carmen provide optimal breeding habitats for this species, favoring its abundance and overcoming the abundance of other mosquito species that need less saline environments for egg eclosion and larval survival, for example, *Ae. aegypti* [72] or species of the *Culex* and *Anopheles* genera [73]. In addition, it has been reported that these species can use freshwater natural habitats such as (epiphytes, cenotes, ponds, and swamps) [40], and artificial habitats (buckets, tires, tanks, and sewage) [44], as well as rainfall accumulation spots for breeding. Moreover, *Ae. taeniorhynchus* has been reported as a migratory species able to move within a range of approximately 30-96 km [74]. Thus, the capacity of females to breed in brackish and freshwater habitats and move long distances contributes to abundance peaks in the coastal and adjacent inland areas of Isla del Carmen and YP during rainfall (June-September) [51] and estuarine flooding seasons (November-February) [27]. *Aedes taeniorhynchus* is implicated in the transmission of several important pathogens that affect humans and animals (Table 1). Therefore, its abundance in Isla del Carmen may pose significant health risks for humans and wildlife populations. Although other less abundant species of *Aedes* were present in Isla del Carmen, it is important to emphasize that they have been implicated in the transmission of pathogens such as viruses or nematodes (Table 1).

It is important to highlight that the species with the highest medical significance in our list was *Ae. aegypti*, the sixth most abundant species, even though our study area is in a semi-conserved area. The presence and abundance of *Ae. Aegypti* could be associated to habitat transformations due to the nearby houses and buildings, and to constant human presence in modified habitat fragments around this zone. It is reported that habitat modifications change the community species composition, and that colonization by synanthrope mosquito species responds to deforestation, loss of native animal and plant species, and to human aggregations [16,47] Moreover, despite of being an urban species, *Ae. aegypti* can survive in heterogeneous landscapes (not fully forested) if it finds the conditions for shelter or breeding [75]. In our study *Ae. Aegypti* was recorded only in NS (75.2%) and RS (24.7%), which could indicate that this species exploits both artificial and natural breeding sites for oviposition. The adaptative plasticity to use breeding sites, such as tree holes, during seasons with higher precipitation has been previously reported in Brazil [76]. Finally, A. aegypti did not show any pattern in the feeding schedule, as it was present in all the sampling hours. It is documented that some factors modifying *Ae. aegypti* flying activity or biting rate are the food availability and artificial light presence, making it active throughout the day, though in lower abundances at night [77,78]. Our results suggest the need for deeper studies on the activity of this important mosquito as a potential vector of pathogenic agents in enzootic cycles.

Earlier studies reported 11 *Psorophora* species in the YP, three of the four species recorded in Isla del Carmen are implicated in the transmission of pathogens such as Ilheus virus (ILHV) [79] and potentially VEE [47]. Since both VEE [80] and ILHV [81] may be distributed within or near the region of Isla del Carmen and involve domestic or wild mammals and birds [82], the presence of numerous migrant bird species [83] that can act as reservoirs might be important factors of risk for human and wildlife populations.

In contrast to the findings of Carpio-Orantes et al. [84], we found evidence of the presence of four *Haemagogus* species in the YP [41,43–45,70,85,86]. Some *Haemagogus* species are involved in the transmission of the sylvatic phase of YFV infection, which has been found mainly in *Hg.* (*Hag.*) *janthinomys* Dyar, 1921, [87], *Hg.* (*Hag.*) *equinus* Theobald, 1903, and *Hg.* (*Hag.*) *lucifer* (Howard, Dyar & Knab 1912), among others [88]. In Isla del Carmen, we recorded only *Hg.* (*Hag.*) *regalis* Dyar & Knab, 1906. Although this species is not related to pathogen transmission, since there are still small populations of wild primates in Isla del Carmen and nearby places, further monitoring of mosquito populations and virologic studies are needed to determine the presence of the wild type YFV and the potential for zoonosis.

The *Culex* genus present in Isla del Carmen included five identified species and two that we cannot identify at the species level using morphological characteristics of females. Mosquito species in this genus are known to be vectors of medical and veterinary relevance. Pathogens such as WNV, St. Louis virus (SLV), ZIK [89–91], and VEE [92] have been found in wild specimens. For example, Baak-Baak et al. [93] determined that *Cx. coronator*, a potential vector of VEE, represents a risk to human health, mainly in rural areas of the YP, including estuarine and mangrove areas similar to those in Isla del Carmen, where people constantly move between villages and the city. Other species in Isla del Carmen, such as *Cx. nigripalpus* Theobald, 1901, have been related to the transmission of WNV and St. Louis encephalitis (SLE) [94]. Thus, as a preventive action, local studies to test for the presence of these pathogens are needed.

Two reported *Coquillettidia* species in the YP were found in Isla del Carmen, *Cq.* (*Rhy.*) *nigricans* (Coquillet, 1904), and *Cq.* (*Rhy.*) *venezuelensis* (Theobald, 1912). Both species were captured during the RS and NS, and no defined activity pattern was observed for any of the species, which were present during daylight and dusk. There is no evidence of medical importance for *Cq. nigricans* [95], but *Cq. venezuelensis* is known to be a natural vector of Flaviviridae, Periburyaviridae, Proxiviridae, and Togaviridae, among other pathogens [96–98], that infect humans, birds, primates and other wild vertebrates.

Among the two *Mansonia* species reported for the YP, only *Ma. dyari* was found in Isla del Carmen. Even though the medical relevance of this species is uncertain, Ortega-Morales and Nava [95] stated that it might be involved in the zoonotic processes of WNV. On the other hand, in Florida, this species was found to be a moderately effective vector for Rift Valley fever virus (RVF) [99], which could be relevant for the rural areas adjacent to Isla del Carmen that are stopover sites for migrant birds, and where cattle and other domestic ruminants are economically important for local inhabitants; however further studies are needed.

Finally, four out of the 12 species of *Anopheles* reported in the YP [43] were found in Isla del Carmen. *Anopheles albimanus* and *An. vestitipennis*, are considered primary malaria vectors in some areas of southern Mexico [100–102], while *An. crucians*, is implicated in the transmission of *Dirofilaria immitis* [103], which infects domestic dogs and cats, cattle [104], and, although rarely, humans [105,106]. So far, *An. gabaldoni*, are not considered important disease vectors, but additional studies are needed.

## Conclusion

This study provides valuable insights into the hourly dynamics, abundance, and diversity of mosquito populations, particularly those of the Culicidae family in the Campeche region, focusing on the Terminos Lagoon area. This research emphasizes the influence of seasonal changes on mosquito populations. Our study revealed that the norte season (NS, characterized by rainfall and cooler temperatures due to cold fronts) exhibited greater mosquito abundance and diversity to that did the dry season (DS) and the rainy season (RS). This pattern is consistent with previous findings, suggesting that mosquito diversity tends to increase during wet seasons. The environmental conditions during the NS allow some species to thrive in large breeding sites with vegetation and standing water, increasing abundance and diversity. Isla del Carmen stands out as a location housing a significant proportion of the mosquito species in the Yucatan Peninsula. The dominance of *Ae. taeniorhynchus* in this area can be attributed to its ability to breed in brackish or freshwater habitats and its migratory nature, which poses potential health risks due to its role in pathogen transmission. This study further highlights the presence of other mosquito species of medical and veterinary relevance, such as those implicated in the transmission of Dengue virus, Ilheus virus, Venezuelan equine encephalitis, West Nile virus, St. Louis virus, Zika virus, and yellow fever virus. This research contributes significantly to understanding mosquito dynamics and the potential health risks associated with specific mosquito species in the Yucatan Peninsula region. Given that there were some undetected species, we emphasize the importance of further monitoring and increased sampling to comprehensively understand mosquito populations in this and other important areas of the coastal southern Gulf of Mexico.

## Acknowledgements

We thank Lourdes Potenciano, Hernán Álvarez and Andrés Reda from ICML-UNAM EL Carmen for supporting the logistics of field work and the sampling sessions.

## Founding

This work did not receive institutional founding.

## Availability of data and materials

Databases on seasonal and hourly species abundance are available on request to the authors.

## Authors’ contribution

JCCD, NVP, and VVS conceived and designed the study. JCCD, NVP, and VVS conducted field surveys, sample processing, and species identification and wrote the manuscript. JCCD, VVS, RPC, and AZJ conducted the statistical analyses of the data. JGCM, OCH, AGP, and MMI revised and approved the final manuscript.

